# Aperiodic EEG and 7T MRSI evidence for maturation of E/I balance supporting the development of working memory through adolescence

**DOI:** 10.1101/2023.09.06.556453

**Authors:** Shane D. McKeon, Maria I. Perica, Ashley C. Parr, Finnegan J. Calabro, Will Foran, Hoby Hetherington, Chan-Hong Moon, Beatriz Luna

## Abstract

Postmortem animal and human models suggest changes through adolescence in aspects of excitatory glutamatergic and inhibitory GABAergic function (E/I) in prefrontal cortex (PFC) suggestive of critical period plasticity at a time of significant cognitive development. Recently, using high field 7T Magnetic Resonance Spectroscopic Imaging (MRSI), we found *in vivo* evidence for increases in PFC glutamate/GABA balance through adolescence into adulthood. We now extend these MRSI findings by investigating, in the same 164 10– 32-year-old participants, its correspondence with EEG aperiodic activity, an independent measure of E/I balance elucidating changes in neural activity. Results showed decreases in PFC aperiodic activity from adolescence to adulthood, that were associated with MRSI measures of glutamate/GABA balance as well as mediating the association between age and EEG aperiodic activity. Further, changes in aperiodic activity predicted performance on a working memory task, indicating a role for E/I based changes in PFC signaling mechanisms in supporting maturation of cognitive control. Taken together, these results suggest that PFC is undergoing critical period plasticity through adolescence evident in both neurotransmitter and neural function that supports cognitive development.

## 1. INTRODUCTION

Prefrontal cortex (PFC) undergoes significant maturation through adolescence including gray matter thinning^1–3^ and changes in task activation^4–7^. In parallel, cognitive control is being refined in adolescence following steep improvements from earlier development^8^. Both PFC maturation and cognitive development become stabilized in adulthood indicating the end of a phase of active specialization that may reflect earlier plasticity^9^. Recently, it has been proposed that adolescence is a developmental critical period for the PFC and its associated functionality^10^. Supporting this hypothesis are studies in rodent models^11,12^ and postmortem human tissue^13,14^ that have shown mechanistic changes consistent with critical period plasticity, including increases in inhibitory, GABAergic parvalbumin (PV) interneurons and decreases in excitatory, glutamatergic processes, resulting in changes in excitation/inhibition (E/I) balance^15^, a hallmark of critical period plasticity^16^. Motivated by these initial findings, we previously used high field 7T MRSI to measure age related changes through adolescence into adulthood in the correspondence of glutamate and GABA function finding evidence of increases in glutamate/GABA balance across frontal cortex^17,18^. These findings correspond with an fMRI GABAergic benzodiazepine challenge study showing developmental increases in E/I balance into adulthood in association cortices including PFC^17,18^. We do not however yet understand the implications of changes in E/I balance at the level of neural activity.

Oscillatory activity has the potential to inform the neural function basis of changes in E/I balance. Synaptic interactions between PV interneurons and pyramidal neurons contribute to the regulation of E/I balance and the production of oscillatory activity, including high-frequency oscillations in the gamma band (30-70 Hz) thought to contribute to cognitive control^19^. Oscillatory activity is measurable through electroencephalogram (EEG), which provides a non-invasive measure of postsynaptic cortical pyramidal neuron function of the brain’s surface. Shifts in oscillatory activity may reflect structural and functional maturation, including the refinement of cortical circuitry, synaptic pruning, myelination, and alterations in E/I circuits^20,21^. However, recent EEG work has demonstrated the importance of accounting for not only the oscillatory periodic activity, but the broadband background aperiodic activity. Aperiodic activity has previously been regarded as “noise” and physiologically irrelevant; however, recent work has shown that this non-cyclical activity reflects neuronal population spiking^22,23^ and may be modulated by task-performance^24,25^. Importantly, it has been found to correspond with states of fluctuating E/I balance in rodent models ^24,26^, and has been shown to be altered in neurological and psychiatric disease states^20,27–30^. This aperiodic activity has been proposed to be measurable by the slope of the power spectral density (PSD) as a function of frequency, that is, how power drops from low to high frequency bands. Previous work has demonstrated this effect by showing that local field potential (LFP) power in neuronal circuits is inversely related to frequency, following a 1/f distribution^20,24,31^ and thought to be representative of the low-pass frequency filtering property of dendrites^31–34^. Furthermore, large scale network mechanisms may also contribute to the 1/f power distribution, as networks with faster time constants, such as excitatory AMPA, have constant power at lower frequencies until they quickly decay, whereas inhibitory GABA currents have slower time constants at higher frequencies and thus delay more slowly as a function of frequency^24,26^, which coupled together, creates the 1/f like slope across broadband frequencies. These results suggest that E/I balance can affect PSD slope: when excitatory activity is greater than inhibitory activity, the 1/f slope will be lower (flatter), whereas it will be larger (steeper) when inhibitory activity is greater than excitatory activity^24^, suggesting the possibility that E/I balance is reflected in the steepness of the PSD slope. The Fitting Oscillations and 1/f (FOOOF) protocol derives the 1/f spectral slope (referred to as the aperiodic exponent) and an offset^24^. The steepness of the slope (i.e., the exponent) has been previously linked to shifts in the E/I balance^24,26–29,35–37^, reductions which reflect shifts toward excitatory activity relative to inhibitory, while increases in the exponent reflect shifts toward inhibitory activity^24,26^. Independently, the PSD offset is thought to reflect broadband shifts in power^24^, reflecting levels of neuronal population spiking^22,38^. Electrocorticogram (ECoG) recordings from macaques have provided empirical support for the FOOOF protocol, showing that sedation by propofol^26^, a general anesthetic that modulates GABA_A_ receptors and effectively decreases the global E/I ratio^39^, resulted in decreases in the power spectral density (PSD) slope^26^. However, there has been little validation of this approach in humans to date, nor direct demonstration of how individual variability in FOOOF measures relates to underlying differences in neurobiology.

Aperiodic activity has also been associated with behavioral performance across a variety of cognitive tasks^24,25,40^, including a reduction in the aperiodic exponent during visuomotor and object recognition tasks relative to baseline^25^, indicating greater predominance of excitatory activity during task as compared to baseline. Further, developmental changes in aperiodic activity have been previously demonstrated, and occur simultaneously with developmental improvements in cognition during adolescence. From early childhood to young adulthood, there is a flattening of the aperiodic exponent and a reduction in offset, potentially reflecting an increase in the E/I balance^20,41–43^. Age-related changes in visual cortical 1/f spectral slopes have also been found to mediate age-related performance in visual working memory tasks for both younger (20-30yo) and older (60-70yo) adults^37^. Given compelling evidence for developmental changes in aperiodic activity, a next step is to understand the other neurobiological indicators of age-related E/I shifts to build models of possible mechanisms underlying maturation.

In this study, we collected a large, multimodal, longitudinal dataset including EEG as well as 7T Magnetic Resonance Spectroscopic Imaging (MRSI). We assessed developmental changes in aperiodic activity in DLPFC as measured by EEG and investigated their association with our initial MRSI findings showing age-related increases in Glu/GABA balance in DLPFC through adolescence^18^ in the same 10–32-year-old participants. We hypothesized that aperiodic activity would decrease through adolescence as Glu/GABA balance increased into adulthood^17^. In line with this hypothesis, we found strong correspondence between decreases in both offset and exponent through adolescence and increasing Glu/GABA balance into adulthood. Finally, we show how EEG- and MRSI-derived measures of E/I balance are associated with behavioral improvements in the latency of working memory responses as measured by the Memory Guided Saccade task. Taken together, this study provides *in vivo* multilevel validation for unprecedented plasticity in PFC through adolescence supporting cognitive development.

## 2. METHODS

### 2.1 Participants

Data was collected on 164 participants (87 assigned female at birth), between 10-32 years of age. This was an accelerated cohort design where participants had up to 3 visits at approximately 18mo intervals, for a total of 286 sessions. Each time point consisted of three visits: a behavioral (in-lab) session, a 7T MRI scan, and an EEG session, typically occurring on different days within 1-2 weeks. Participants were recruited from the greater Pittsburgh area and were excluded if they had a history of loss of consciousness due to a head injury, non-correctable vision problems, learning disabilities, a history of substance abuse, or a history of major psychiatric or neurologic conditions in themselves or a first-degree relative. Patients were also excluded if any MRI contradictions were reported, including but not limited to, non-removable metal in their body. Participants or the parents of minors gave informed consent with those under 18 years of age providing assent. Participants received payment for their participation. All experimental procedures were approved by the University of Pittsburgh Institutional Review Board and complied with the Code of Ethics of the World Medical Association (Declaration of Helsinki, 1964).

### 2.2 Data Acquisition and Preprocessing

#### 2.2.1 Electrophysiological (EEG) Data

Concurrent EOG (electrooculogram) and high-impedance EEG was recorded using a Biosemi ActiveTwo 64-channel EEG system located in the PWPIC Psychophysiology Laboratory. EEG sessions were conducted in an electromagnetically shielded room while stimuli were presented by a computer approximately 80 cm away from participants. Resting state data was collected from four eyes open and eyes closed conditions, each lasting one minute and alternating between conditions. Initial data was sampled at 1024 Hz and resampled at 150 Hz during preprocessing. Scalp electrodes were referenced to external electrodes corresponding to the mastoids due to its proximity to the scalp and low signal recording. An initial bandpass filter was set to 0.5 – 90 Hz. Data were preprocessed using a revised processing pipeline compatible with EEGLAB^44^, which removed flatline channels (maximum tolerated flatline duration: 8 seconds), low-frequency drifts, noisy channels (defined as more than 5 standard deviations from the average channel signal), short spontaneous bursts, and incomplete segments of data. Deleted channels were replaced with interpolated data from surrounding electrodes. The resulting data was referenced to the average reference. As a final preprocessing step, independent component analysis (ICA) was performed to identify eye-blink artifacts and remove their contribution to the data.

#### 2.2.2 Magnetic Resonance Spectroscopy

MRSI methods have been previously reported in Perica et al. (2022)^18^. Briefly, data were acquired at the University of Pittsburgh Medical Center Magnetic Resonance Research Center using a Siemens 7T scanner. Structural images were acquired using an MP2RAGE sequence (1 mm isotropic resolution, TR/TE/flip angle 1/ flip angle 2: 6000 ms/2.47 ms/4^0^/5^0^). MRSI including GABA and glutamate were acquired using a J-refocused spectroscopic imaging sequence (TE/TR = 2 ×17/1500 ms) and water suppression was performed using a broad band semi-selective refocusing pulse and frequency selective inversion recovery^45^. Radiofrequency (RF) based outer volume suppression was used to minimize interference in the signal from extracerebral tissue ^46^. An 8 × 2 1 H transceiver array using 8 independent RF channels was used to acquire data. High order shimming was used to optimize homogeneity of the B_0_ magnetic field. The 2D CSI oblique axial slice was acquired with conventional rectangular phase encoding of 24 × 24 over a FOV of 216 × 216 mm (10 mm thick, 0.9 ×0.9 ×1.0 cm nominal resolution), and was positioned to include Brodmann Area 9 and pass through the thalamus.

#### 2.2.3 Memory Guided Saccade Task

Participants performed a memory guided saccade (MGS) task to assess working memory (see Figure 1) during the EEG session. The trial began with fixation to a blue cross for 1 sec. The participant was then presented with a peripheral cue with a surrounding scene picture to be used in a different study, in an unknown location along the horizontal midline (12.5 or 22.2 degrees from central fixation to left or right of center), where they performed a visually guided saccade (VGS) to the target and maintained fixation. Once the cue disappeared, the participant returned their gaze to the central fixation point and fixated for a variable delay epoch (6-10 sec) during which they were to maintain the location of the peripheral target in working memory. Once the central fixation disappeared the participant performed a memory guided saccade to the recalled location of the previous target. The trial ended when participants were presented with a white fixation cross that served as the ITI (1.5 - 15sec). Participants performed 3 runs of the MGS task, each containing 20 trials.

**Figure 1.**
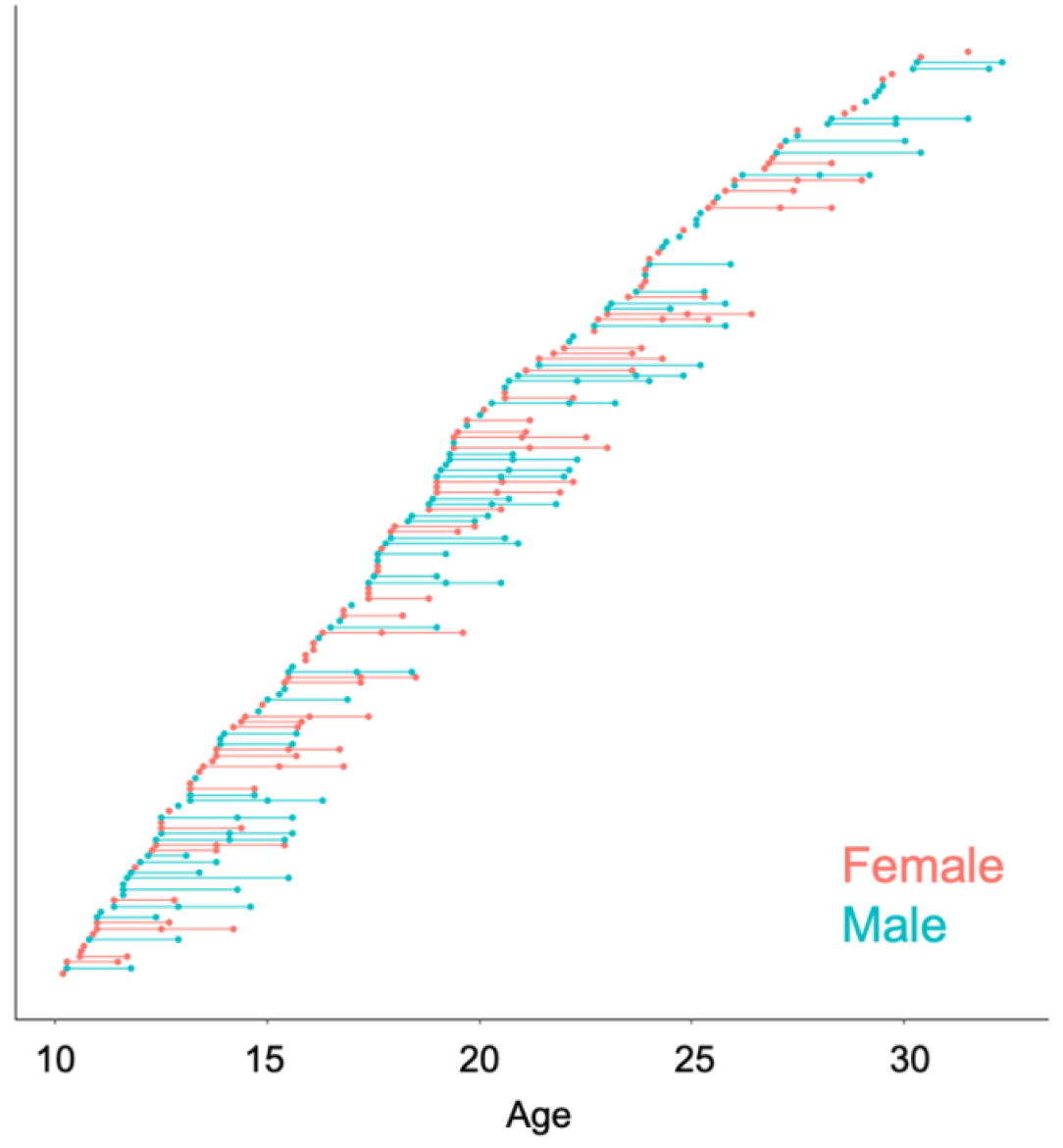
Distribution of participants, depicting the number of visits per participant: each point represents an individual scan, with lines connecting individual participants

**Figure 2.**
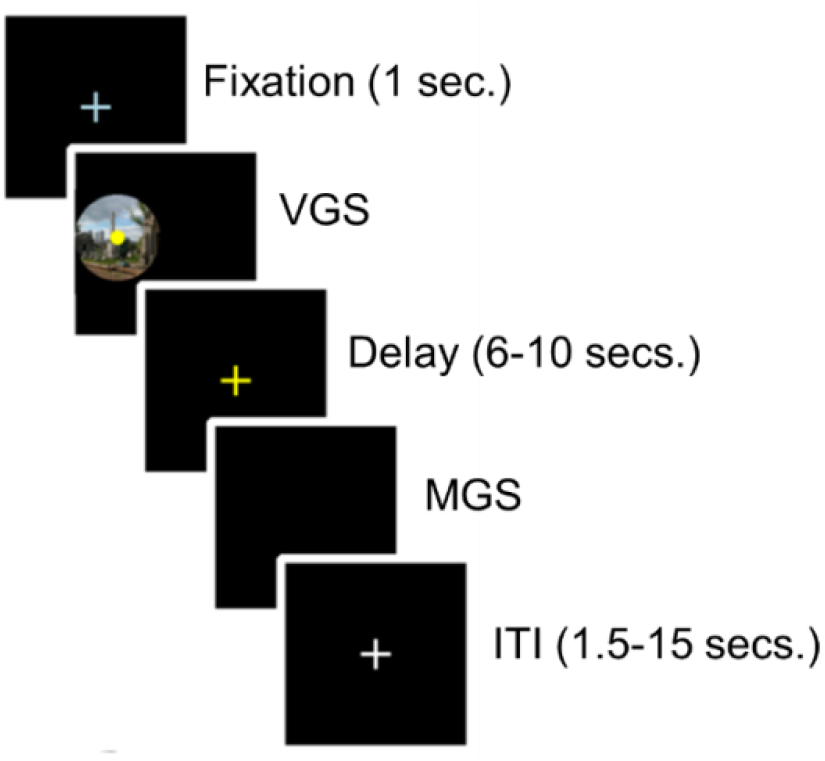
Memory Guided Saccade Task. Epochs from top to bottom: Participants were presented with a fixation cross. Once fixation was extinguished, participants performed a saccade to a peripheral dot stimulus on top of a scene (the scene is intended for an incidental memory task for a separate aim of this project). After the peripheral target was extinguished, participants were presented with a yellow fixation cross while remembering the location of the previous target. When the fixation cross was extinguished, participants performed a second saccade to the remembered target location. A variable ITI (1.5-15 sec) with a fixation white cross occurred between trials.

Task performance was assessed based on horizontal electrooculogram (hEOG) channels recorded from facial muscles (see acquisition details below). At the start of the session, participants performed a calibration procedure in which they fixated a series of 20 dots sequentially, positioned along the horizontal midline and spanning the width of the screen. These were used to generate a calibration curve relating hEOG voltage to horizontal screen position. Eye position during the MGS task was used to derive output measures using this calibration data by aligning hEOG signals to different stimulus triggers. These were used to calculate VGS & MGS response latencies, as the time difference between the beginning of the recall epoch and the initiation of the VGS and MGS eye movements respectively, and saccadic accuracy, measured as the closest stable fixation point during the recall period to the fixated location during the initial visually guided fixation epoch.

#### 2.2.4 Spatial Span Task

Participants preformed the Spatial Span Test from the Cambridge Neuropsychological Test Automated Battery (CANTAB)^47^ that assesses working memory capacity. During the task white squares appear on the screen, some of which will briefly change color in a variable sequence. The participant must select the boxes that changed color in the same order that they were presented. The number of boxes in the sequences begins with two and increases, after each successful trial, to a maximum of nine. The outcome measure was the maximum sequence length the participant successful completed, between 2 and 9.

### 2.3 Data Analysis

#### 2.3.1 EEG Analyses

Power spectral density (PSD) was calculated separately for each participant and electrode, corresponding to the left and right DLPFC (Right: F4, F6, F8; Left: F3, F5, F7), across the continuous resting state EEG using Welch’s method implemented in MATLAB (2s Hamming window, 50% overlap). The Fitting Oscillations and One Over *f* (FOOOF) python toolbox (version 1.0.0; https://fooof-tools.github.io/fooof/) was used to characterize the PSD as a combination of an aperiodic component with overlying period components, or oscillations^24^ (see Fig 3A). Oscillations were characterized as frequency regions with power above the aperiodic component, modeled as a Gaussian, and are referred to as ‘peaks’. The aperiodic component *L*(*f*) was extracted from the 1-50Hz frequency range of each power spectrum (*aperiodic_mode* = ‘fixed’, *peak_width_limits* = [0.5, 12], *min_peak_height* = 0, *peak_threshold* = 2, *max_n_peaks* = 4, default settings otherwise), and is expressed as:

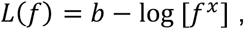

with a constant offset *b* and the aperiodic exponent *x*. When the PSD is plotted on a log-log axis, the constant offset *b* refers to the y-intercept and the aperiodic exponent *x* corresponds to the slope of a line (see Figure 3). We used the ‘fixed’ setting as we did not expect a “knee” in the power spectrum. This assumption was supported upon visual inspection of each PSD. The average *R*^*2*^ of the spectral fits was 0.98 for the eyes closed condition and 0.97 for the eyes open condition, reflecting good fits.

**Figure 3.**
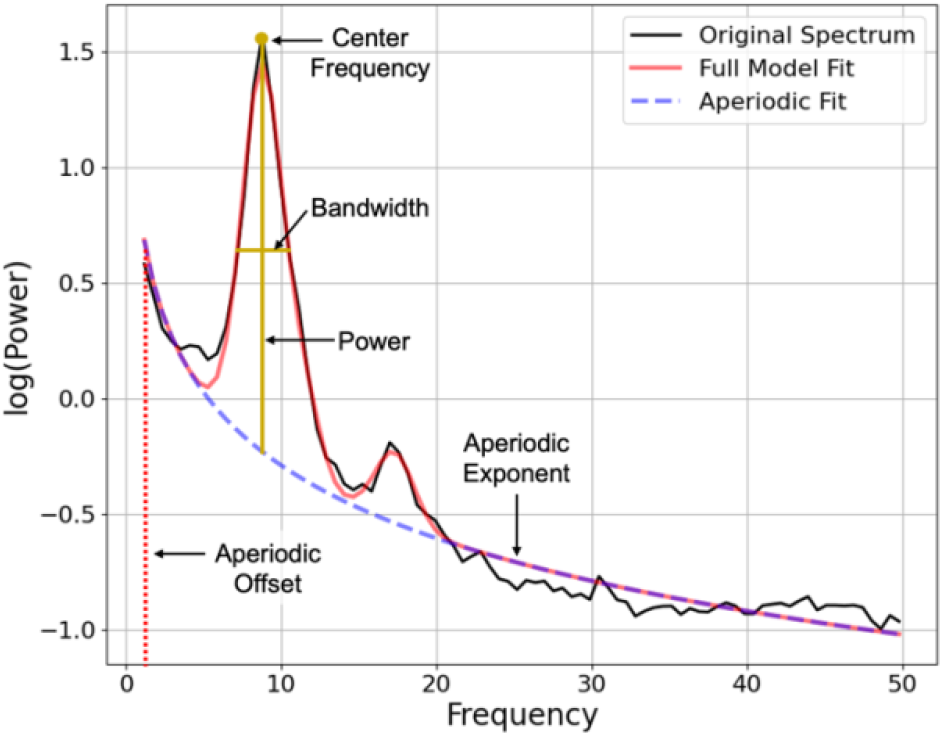
Example FOOOF model fit from a representative participant during eyes closed resting state displaying the aperiodic offset (dashed red) and the aperiodic slope (i.e., the exponent) dashed purple) across the entire frequency range (1-50 Hz). The center frequency, power, and bandwidth are highlighted for the peak present around 10 Hz.

#### 2.3.2 MRS Analyses

Analysis details have been previously reported in Perica et al. (2022)^18^. LCModel was used to obtain estimates of individual metabolites^48^, including macromolecule components and 14 basis metabolite functions (N-acetylaspartate (NAA), N-acetylaspartylglutamate (NAAG), aspartate, lactate, creatine (Cre), γ-aminobutyric acid (GABA), glucose, glutamate (Glu), glutamine, glutathione, glycerophosphorylcholine (GPC), phosphorylcholine (PCh), myoinositol, and taurine), of which we focus on GABA and Glu for the purposes of this study.

A 2D CSI oblique axial slice was acquired, as outlined in Perica et al. (2022), to ensure that the dorsolateral prefrontal cortex (DLPFC) was included and angled to pass through the thalamus. Regions of interest included right and left DLPFC due to its role in working memory. Data quality exclusion was performed at the ROI level. All spectra and model fit outputs were visually inspected and used as first-pass exclusion criteria, followed by Cramer-Rao Lower Bound (CRLB) > 10 for the three major metabolite peaks (GPC/Cho, NAA/NAAG, and Cr), and CRLB > 20 for glutamate and GABA. All glutamate and GABA levels are given as ratios to creatine (Cr). Metabolite levels were corrected for fractional gray matter within the voxel as well as date to control for any changes in scanner precision over time, while controlling for age. Glu/GABA ratio was calculated by creating a ratio of Glu/Cr to GABA/Cr. Glu-GABA asymmetry was calculated by taking the absolute value residual of the linear model of the association between Glu/Cr and GABA/Cr. A threshold of 2 standard deviations from the mean was used to exclude statistical outliers.

#### 2.3.3 Behavioral Analyses

To examine age-related effects in behavioral measures, including accuracy and response latency, generalized additive mixed models (GAMMs) were implemented using the R package mgcv^49^. Preliminary outlier detection was conducted on a trial level basis. Express saccades of less than 100ms, believed to be primarily driven by subcortical systems^50^, were excluded. Position error measures greater than 23 degrees from the target were excluded as implausible since they exceeded the width of the screen. The remaining trials for each participant were combined, creating average performance measures and trial-by-trial variability measures for each participant. Finally, for group level analyses, outlier detection was performed, excluding participants more than +/-2 SDs away from the mean. A separate model was performed for each of the behavioral measurements: MGS accuracy, MGS latency, as well as the trial-by-trial variability of each measure. To correct for multiple comparisons between the four behavioral measures, Bonferroni correction was employed which resulted in a corrected alpha of .0125 (*p* = .05/4 = .0125). For all behavioral measures showing significant age-related effects, we performed analyses to identify specific periods of significant age-related differences. To do so, a posterior simulation was performed on the first derivative of the GAM model fits. Consistent with previous work^49,51,52^, 10000 simulated GAM fits and their derivatives (generated at age intervals of 0.1 years) were computed from a multivariate normal distribution; vector means and covariance of which corresponded to the fitted GAM parameters. Confidence intervals (95%) were generated from the resulting derivatives. The periods of age-related growth were derived from the ages corresponding to when the confidence interval did not include zero (p < 0.05).

For the spatial span task, we used generalized additive mixed models (GAMMs), implemented using the R package mgcv^49^, to assess age-related effects in maximum sequence length. We then performed analysis to identify specific periods of significant age-related differences. To do so, a posterior simulation was performed on the first derivative of the GAM model fits. Consistent with previous work^49,51,52^, 10000 simulated GAM fits and their derivatives (generated at age intervals of 0.1 years) were computed from a multivariate normal distribution; vector means and covariance of which corresponded to the fitted GAM parameters. Confidence intervals (95%) were generated from the resulting derivatives. The periods of age-related growth were derived from the ages corresponding to when the confidence interval did not include zero (p < 0.05).

### 2.4 Statistical Analyses

#### 2.4.1 Characterizing age-related trends in 1/f aperiodic activity

To assess developmental trajectories of aperiodic activity, we implemented GAMMs on aperiodic parameter (exponent and offset), including random intercepts estimated for each participant. Regression splines were implemented (4 degrees of freedom) to assess linear and non-linear effects^49,53^. We first tested for a main effect of age on aperiodic parameter while controlling for hemisphere (either ‘right’ or ‘left’ DLPFC) and condition (eyes open or eyes closed during resting state), seen in Model 1 in Supplement. We additionally tested for age-by-hemisphere interactions while controlling for condition (Supplement Model 2), and age-by-condition interactions while controlling for region (Supplement Model 3). Correlations between the exponent and the offset, for both the eyes open and eyes closed conditions were calculated using Pearson correlations.

#### 2.4.2 Characterizing age-related trends in MRSI-derived metabolite levels

To assess age-related change in the Glu/GABA ratio and the Glu-GABA asymmetry in right and left DLPFC, we used GAMM models, including random intercepts estimated for each participant. Regression splines were implemented (4 degrees of freedom) to assess linear and non-linear effects^49,53^. We first tested for a main effect of age on the MRS measure while controlling for hemisphere (either ‘right’ or ‘left’ DLPFC), and perfect grey matter from the MRI voxel (Supplement Model 4). We additionally tested for age-by-hemisphere interactions while controlling for fraction of grey matter (Supplement Model 5). For further analysis, fraction of gray matter in the voxel was residualized out of MRSI estimates to control for the effect of gray matter.

#### 2.4.3 Characterizing the relationship between 1/f aperiodic activity and MRS

We next investigated the relationship between the individual aperiodic parameters (exponent and offset) on MRSI-derived measures of the Glu-GABA asymmetry using linear mixed effect models (lmer function, lme4 package in Rstudio^54^). Using AIC, we determined an inverse age (age^-1^) model was more appropriate for both the exponent vs MRS and offset vs MRS associations (Supplemental Table 1), thus each model controlled for inverse age. We first tested for significant main effects of the MRS measure on the aperiodic parameter while controlling for age^-1^, condition (eyes open or eyes closed), and hemisphere (left or right DLPFC) (Supplement Model 6). We additionally tested for MRS-by-age^-1^ interactions while controlling for hemisphere and condition (Supplement Model 7). To account for the four aperiodic measures, Bonferroni correction was used for multiple comparisons at p_bonferroni_ = 0.012.

**Table 1.** Linear Mixed Effect Models examining associations between EEG-derived measures of aperiodic activity and MRS-derived measures of glutamate (Glu), GABA, and the Glu GABA imbalance measure. (* p < 0.01; ** p < 0.001; *** p < 0.0001)

|  |  | Main effect of MRS |  |  | MRS*inverse age interaction |  |  | Main effect of Hemisphere |  |  | Main effect of Condition |  |  |
| --- | --- | --- | --- | --- | --- | --- | --- | --- | --- | --- | --- | --- | --- |
| | | $\beta$ | t | p | $\beta$ | t | p | $\beta$ | t | p | $\beta$ | t | p |
| Exponent | Glu | 0.07 | 2.30 | **0.04** | -3.11 | -1.88 | 0.12 | 0.01 | 1.03 | 0.61 | -0.15 | -14.16 | **0.00** |
|  | GABA | 0.05 | 0.88 | 0.76 | 1.90 | 0.54 | 1.00 | 0.01 | 0.47 | 1.00 | -0.15 | -13.58 | **0.00** |
|  | Asymmetry | 0.12 | 2.22 | **0.05** | 0.71 | 0.23 | 1.00 | 0.01 | 0.43 | 1.00 | -0.15 | -13.44 | **0.00** |
| Offset | Glu | 0.08 | 2.62 | **0.02** | -3.25 | -1.90 | 0.11 | 0.05 | 4.34 | **0.00** | -0.15 | -14.15 | **0.00** |
|  | GABA | 0.08 | 1.23 | 0.44 | -0.51 | -0.13 | 1.00 | 0.04 | 3.10 | **0.00** | -0.16 | -13.66 | **0.00** |
|  | Asymmetry | 0.09 | 1.53 | 0.26 | -0.20 | -0.06 | 1.00 | 0.04 | 3.40 | 0.00 | -0.15 | -13.64 | **0.00** |

#### 2.4.4 Characterizing the relationship between 1/f aperiodic activity and behavior

To assess associations between 1/f aperiodic parameters (exponent and offset), MGS behavioral measures (accuracy, accuracy variability, response latency, and response latency variability), and spatial span measure (maximum length of sequence), we used linear mixed effect models (lmer function, lme4 package in Rstudio^54^). Using AIC, we determined an inverse age (age^-1^) model was more appropriate for both the exponent vs behavior and offset vs behavior associations (Supplemental Table 2). We first tested for significant main effects of the behavioral measure on the aperiodic parameter while controlling for inverse age (age^-1^), condition (eyes open or eyes closed), and hemisphere (left of right DLPFC) (Supplement Model 8). We additionally tested for behavior-by-inverse age interactions while controlling for condition and hemisphere (Supplement Model 9). Bonferroni correction was used for multiple comparisons.

#### 2.4.5 Characterizing the relationship between MRS and behavior

To assess associations between MRS measures (asymmetry) and MGS behavioral measures (accuracy, accuracy variability, response latency, and response latency variability), and spatial span measure (maximum length of sequence), we used linear mixed effect models (lmer function, lme4 package in Rstudio^54^). Using AIC, we determined an inverse age model was best for the glutamate, GABA, and asymmetry vs behavioral associations (Supplemental Table 2). We first tested for significant main effects of the behavioral measure on the aperiodic parameter while controlling for age^-1^ and hemisphere (left of right DLPFC) (Supplement Model 10). We additionally tested for behavior-by-age interactions while controlling hemisphere (Supplement Model 11). Bonferroni correction was used for multiple comparisons.

## 3. RESULTS

### 3.1 Working Memory Performance Improves across Adolescence

Consistent with our prior findings using the MGS task^50,55,56^, behavioral performance improved with age for all MGS metrics including increased accuracy (*F* = 38.19, *p* = <2e-16; Figure 4A), decreased response latency (*F* = 17.91, *p* = <2e-16; Figure 4B), and decreased trial-to-trial variability in both accuracy (*F* = 29.1, *p* = <2e-16; Figure 4C) and response latency (*F* = 52.04, *p* = <2e-16; Figure 4D). Significant developmental improvements were found throughout adolescence (11-24 years of age) for MGS accuracy and between 10-24 years of age for trial-by-trial variability for MGS accuracy (Figure 4A and 4C). MGS latency showed significant increases in adolescence (12-17 and 19-22 years of age, Figure 4B), while decreases in MGS variability in latency continued into the twenties (Figure 4D). Furthermore, working memory capacity also improved across adolescence via the Spatial Span task as seen in a significant increase in the maximum number of boxes presented in the sequence (*F* = 11.35, *p =* 0.00035; Figure 5). Significant developmental improvements were found between 10-17 years of age.

**Figure 4.**
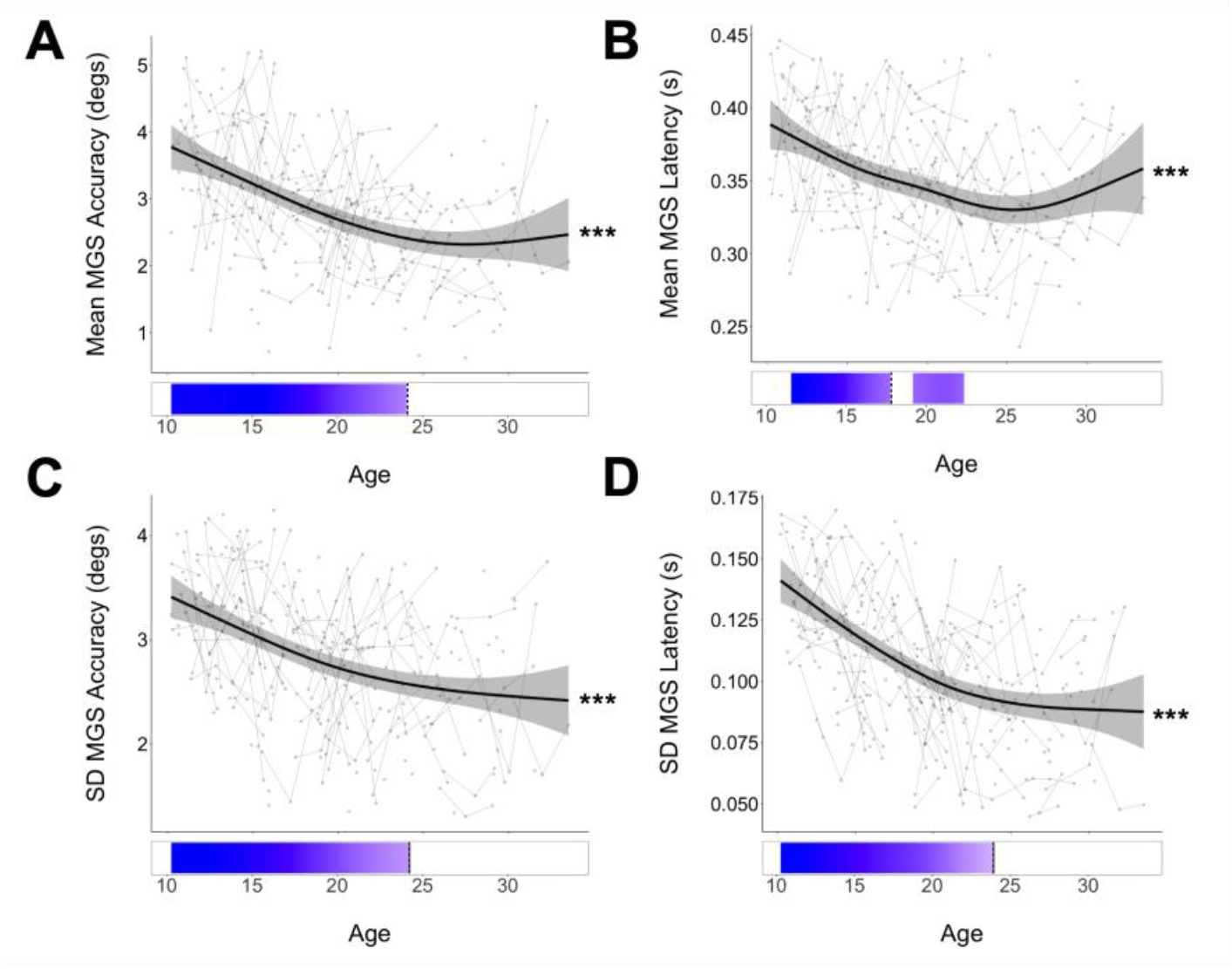
**A**. Mean MGS accuracy (degs.). Growth rate analyses show significant periods of improvement between 11-24 years of age **B**. Mean MGS response latency (s). Growth rate analyses show significant periods of improvement between 12-17 and 19-22 years of age **C**. Trial-by-trial variability in MGS accuracy (degs.). Growth rate analyses show significant periods of improvement between 10-24 years of age **D**. Trial-by-trial variability of MGS response latency. Growth rate analyses for show significant periods of improvement between 10-24 years of age. Blue bars indicate regions of significant age-related change in MGS performance based on the derivative of the GAM model fit. (* p < 0.01; ** p < 0.001; *** p < 0.0001)

**Figure 5.**
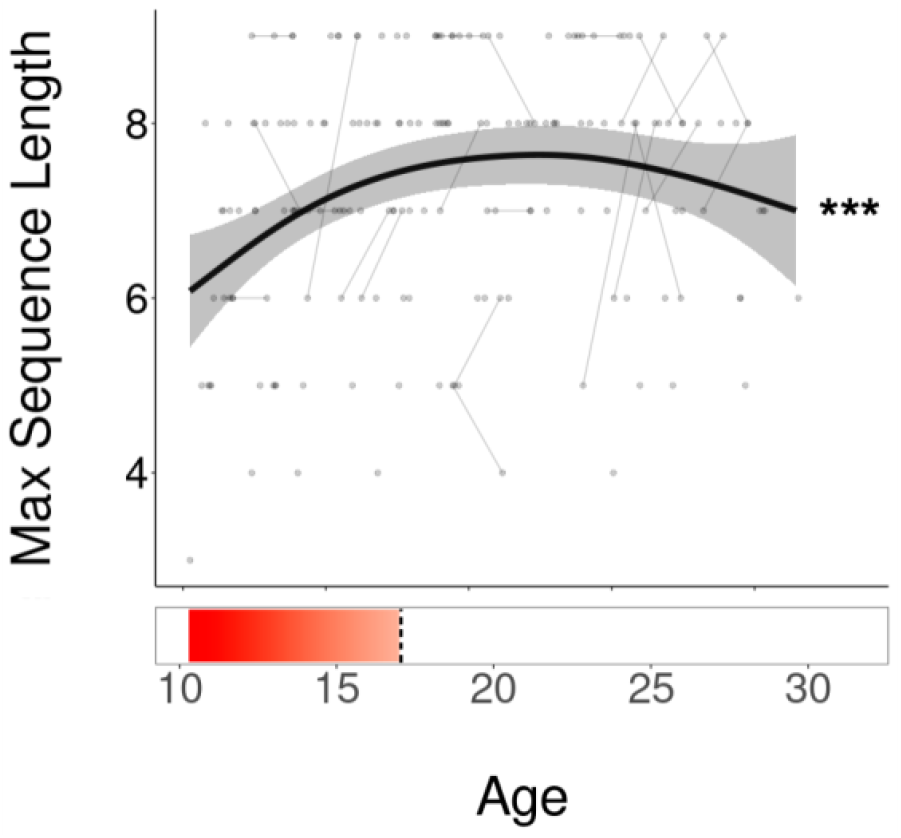
Maximum number of boxes presented to the participant vs age. Growth rate analyses show significant periods of improvement between 10-17 years of age. (*** p < 0.0001)

### 3.2 Glutamate GABA Asymmetry Decreases through Adolescence

As in our prior cross-sectional study (Perica et al, 2022), glutamate showed age-related decreases across adolescence (F = 5.72, *p* = 0.02; Figure 6A), while there were no significant age effects with GABA (F = 0.04, *p* = 0.84; Figure 6B). As previously, we assessed Glu/GABA balance by assessing asymmetry, an individual-level measure of the correlation between glutamate and GABA which was defined as the absolute difference in age-adjusted Glu and GABA levels (see methods), was found to significantly decrease with age (F = 13.60, *p* = 0.0002; Figure 6D). As expected, due to greater inter-subject variability at younger ages in Glu and GABA undermining the ability to assess age related changes in balance^18^, the ratio was not associated with age (F = 3.68, *p* = 0.10; Figure 6C). Thus, analyses going forward utilized the Glu/GABA asymmetry measure. Additional statistics regarding main effects of hemisphere and age-by-hemisphere interactions can be found in the supplement (Figure S1).

**Figure 6.**
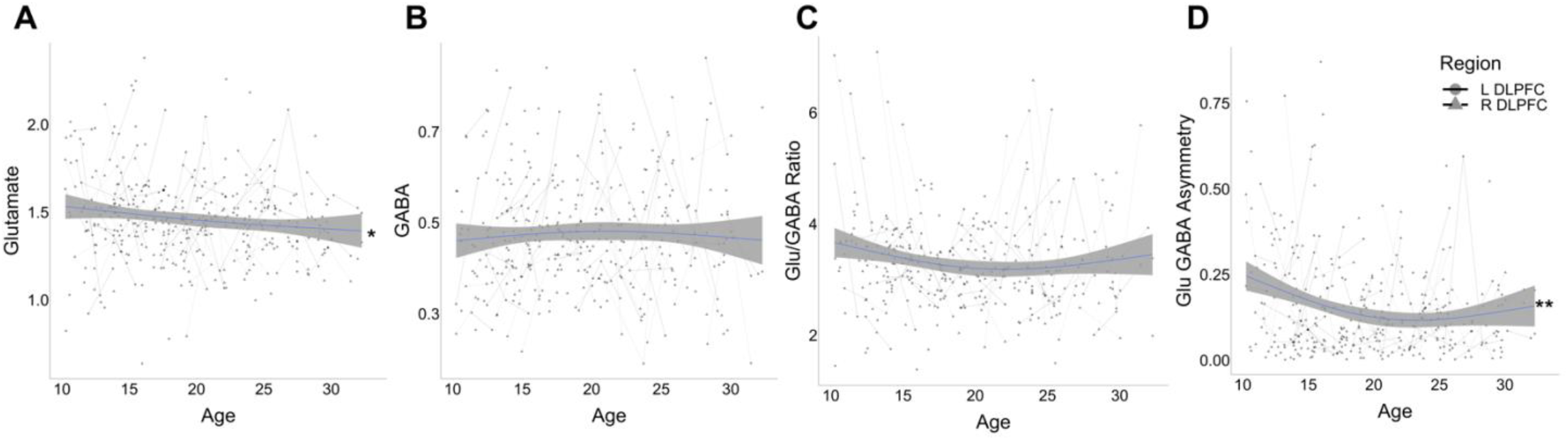
MRSI derived metabolite levels across adolescence. **A**. Glutamate vs age across adolescence. **B**. GABA vs age across adolescence **C**. Glu/GABA Ratio vs age across adolescence. **D**. Glutamate GABA asymmetry vs age. (* p < 0.01; ** p < 0.001; *** p < 0.0001)

### 3.3 1/f Aperiodic Activity Decreases through Adolescence

To assess changes in signal processing characteristics through adolescence, each FOOOF model parameter was tested for associations with age. For the aperiodic EEG exponent measure, which captures the steepness of the relationship between power and frequency, we found significant age-related decreases (F = 61.85, *p* < 0.0001; Figure 7A). The aperiodic offset, which captures the overall power independent of frequency, was also found to significantly decrease with age (F = 237.1, p < 0.0001; Bottom panel in Figure 7B). The exponent and offset were highly correlated across both conditions (*ρ* = 0.75, *p* < 2.2e-16) (Figure 7C). Additional statistics including the effect of condition (eyes open vs eyes closed) and hemisphere (right vs left DLPFC) can be found in the supplemental Figure 2.

**Figure 7.**
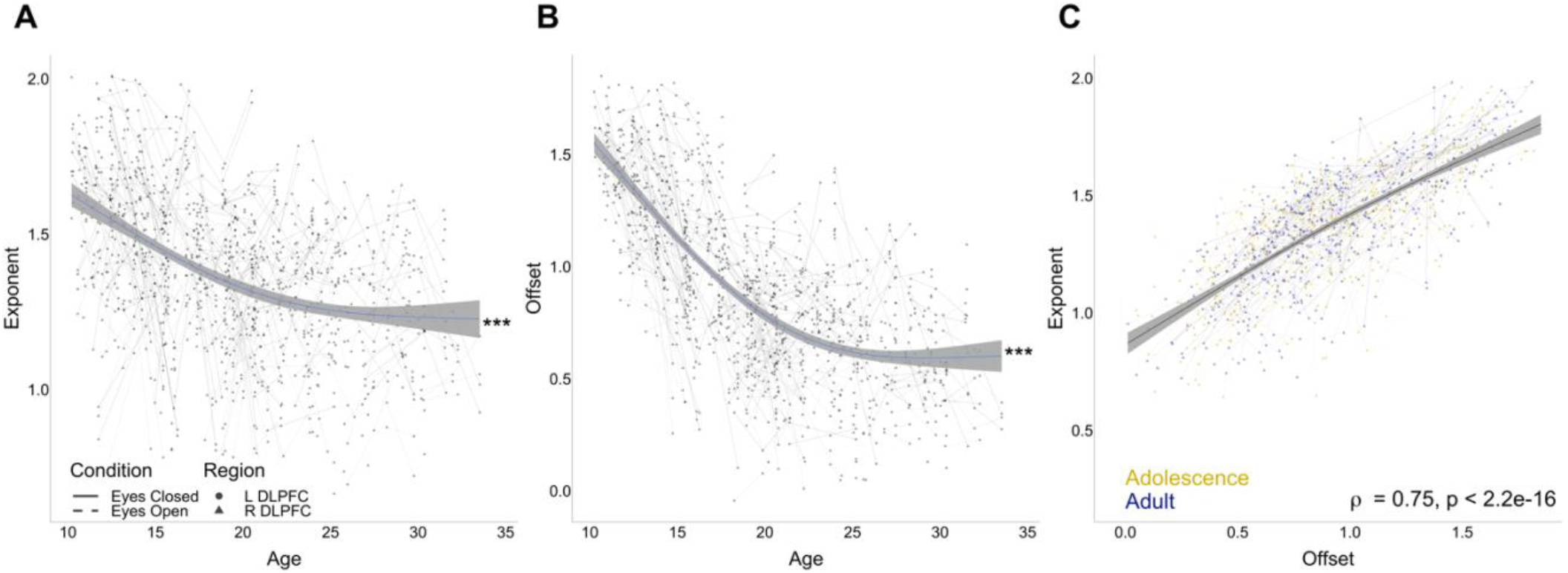
Aperiodic activity of EEG during resting state. **A**. Exponent across adolescence across both eyes closed and eyes open conditions. **B**. Offset across adolescence across both eyes closed and eyes open. **C**. Correlation between exponent and offset values for both the left and right DLPFC for both conditions.

### 3.4 Associations between 1/f Aperiodic Activity and MRS

To characterize the functional associations of MRSI-derived metabolite levels, we next assessed associations between EEG-derived FOOOF measures and Glu-GABA asymmetry. All statistics were conducted controlling for inverse age and are reported in Table 1 with corrected p values. The Glu-GABA asymmetry significantly increased with increasing aperiodic exponent (β = 0.12, t = 2.22, *p* = 0.05; Figure 8A) but not offset (β = 0.09, t = 1.53, *p* = 0.26; Figure 8B). To interrogate whether the effects of the glutamate-GABA asymmetry on the aperiodic activity was driven by glutamate or GABA, we conducted post hoc analyses of these associations. Here, we found glutamate to significantly increase with increasing exponent (β = 0.07, t = 2.30, *p* = 0.04; Figure 8C), and increasing offset (β = 0.08, t = 2.62, *p* = 0.02; Figure 8D). GABA did not have a significant association with either exponent (β = 0.05, t = 0.88, *p* = 0.76; Figure 8E) nor offset (β = 0.08, t = 1.23, *p* = 0.44; Figure 8F).

**Figure 8.**
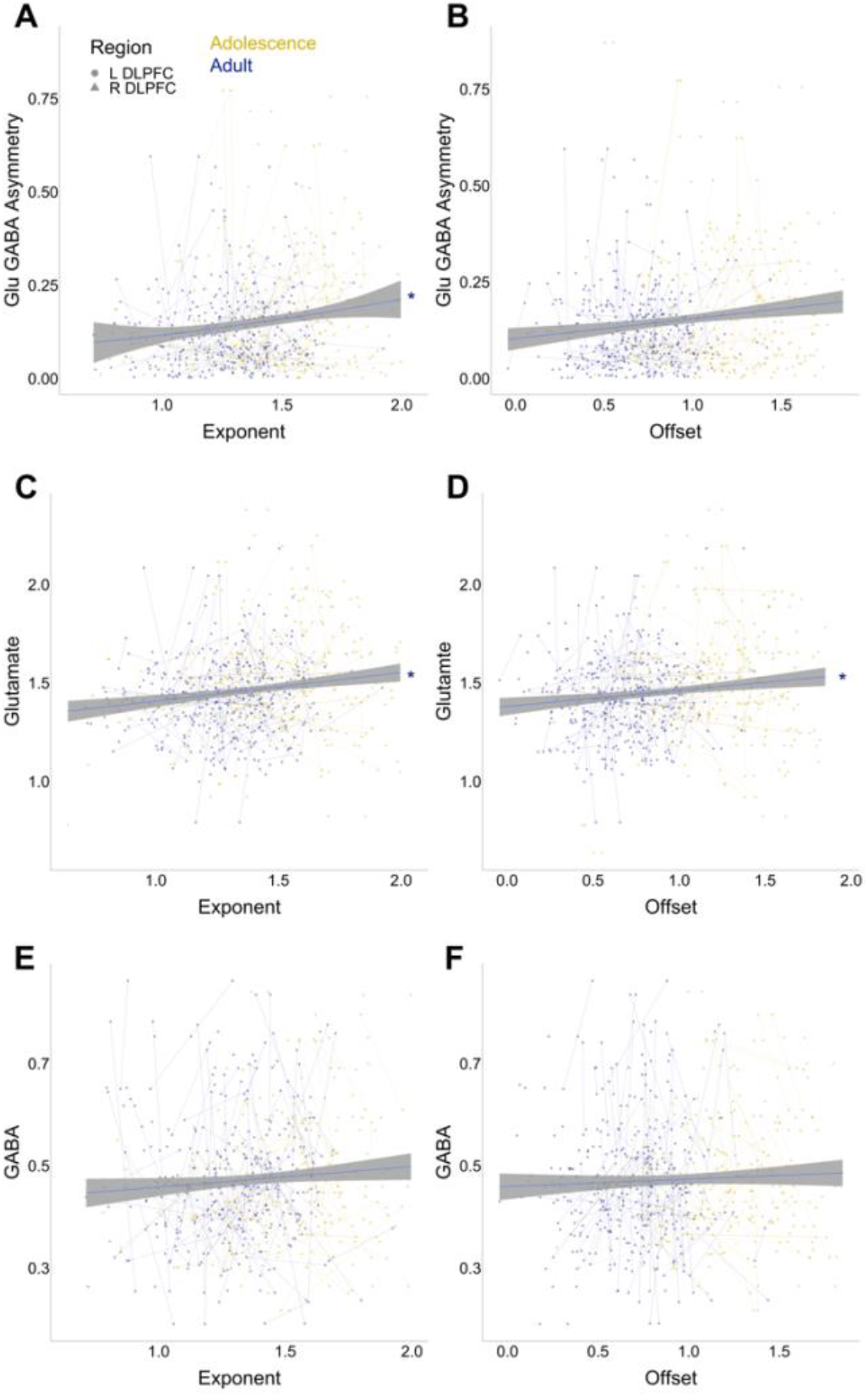
Aperiodic EEG activity associations with MRS measures. Adolescence is all participants under 18 (gold) and adult represents all participants 18 and above (purple). **A**. Exponent vs Glu GABA asymmetry. **B**. Offset vs Glu GABA asymmetry **C**. Exponent vs Glutamate **D**. Offset vs Glutamate **E**. Exponent vs GABA **F**. Offset vs GABA

To further interrogate the contribution of metabolite levels to the generation of aperiodic activity, we performed mediation analyses for MRSI measures that showed significant associations with aperiodic EEG. Glutamate was not a significant mediator of age-related changes in offset (ACME: −0.00045, 95% CI [-0.0011, 0.00], *p* = 0.056; Figure 9A). However, it did significantly mediate age-related changes in exponent (ACME: - 0.00048, 95% CI [-0.0012, 0.00], *p* = 0.028*; Figure 9B). We found that the Glu-GABA asymmetry was a significant mediator of age-related changes in the FOOOF exponent parameter (ACME: −0.00053, 95% CI [- 0.0012, 0.00], *p* = 0.04; Figure 9C).

**Figure 9.**
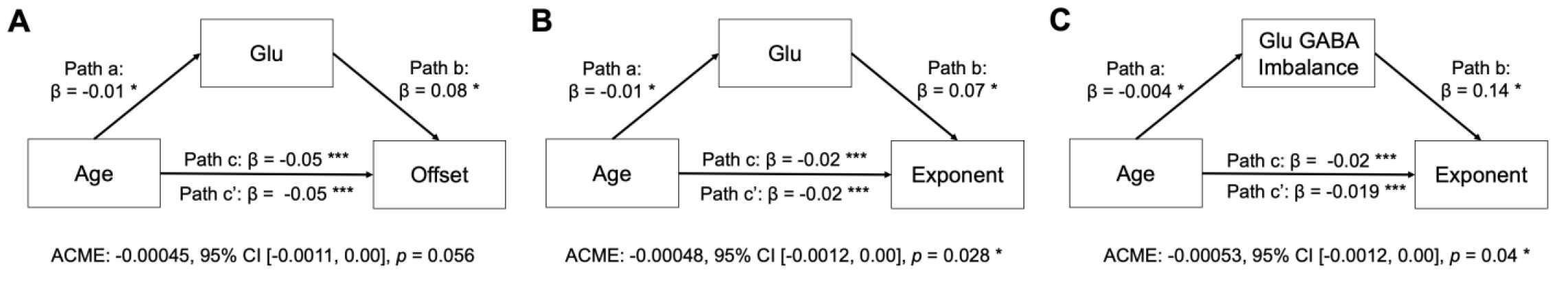
Mediation Analyses. **A**. Glutamate mediation on age-related changes on offset **B**. Glutamate mediation on age-related changes on exponent **C**. Glu GABA asymmetry mediation on age-related exponent.

### 3.5 Increased Aperiodic Activity is associated with Improved Working Memory Performance

To characterize the functional significance of our age-related changes in 1/f aperiodic activity, we assessed their relationship with our MGS working memory task, and the spatial span task. All associations and their statistics can be found in Supplement Table 3 and Figure S1. Exponent showed a trend-level positive association with MGS response latency (*β* = −0.63, t = −2.44 *p* = 0.06; Figure 10A), and offset was significantly associated with increased response latency (*β* = −1.04, t = −3.60, *p* = 0.001; Figure 10B). Furthermore, lower exponent (*β* = −0.04, t = −3.09, *p* = 0.002; Figure 10C) and offset (*β* = −0.04, t = −3.12, *p* = 0.002; Figure 10D) were associated with increased maximum sequence lengths indicating improved spatial span performance.

**Figure 10.**
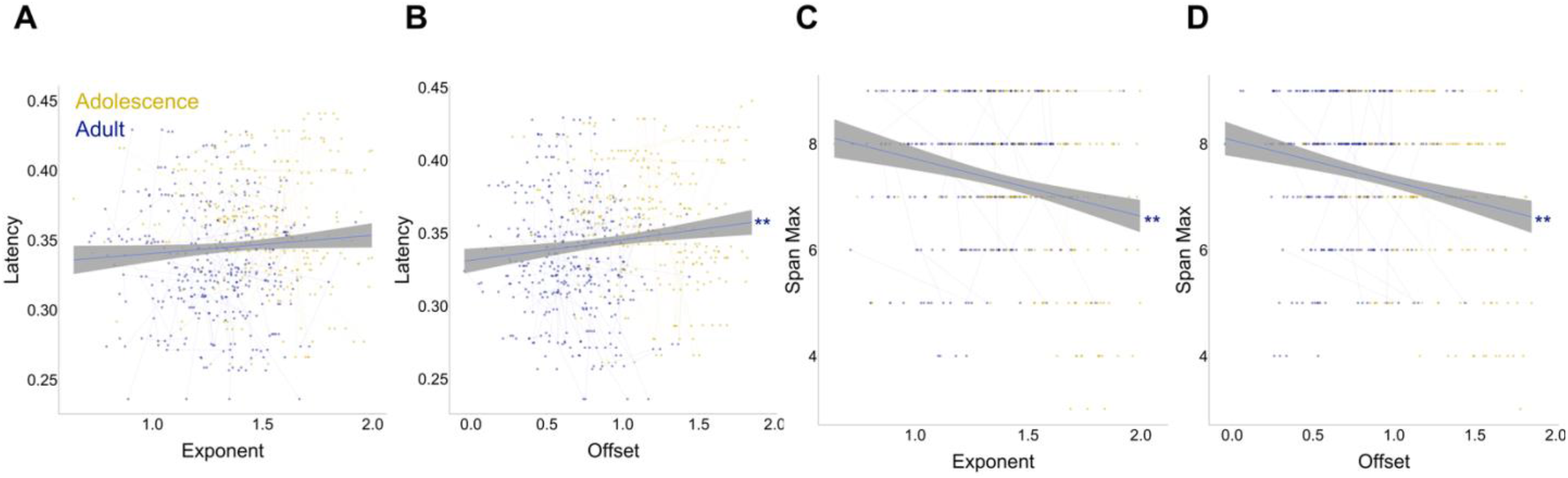
Aperiodic EEG activity associations with behavioral measures. Adolescence is all participants under 18 (gold) and adult represents all participants 18 and above (purple). **A**. Exponent vs MGS response latency **B**. Offset vs MGS response latency **C**. Exponent vs Spatial span max sequence length **D**. Offset vs Spatial span max sequence length. (** p < 0.001)

To characterize the functional significance of our age-related changes in our MRS measures, we assessed their relationship with our MGS working memory task and the spatial span task. All associations and their statistics can be found in Supplement Table 3 and Figure S3. Glu GABA Asymmetry had no relationship with response latency (*β* = −0.06, t = −0.31, *p* = 1; Figure 11A). There was a significant latency-by-inverse age interaction on Glu GABA Asymmetry (*β* = 32.33, t = 3.01, *p* = 0.01). To interrogate whether the effects of the trending Glu GABA Asymmetry on the behavior was driven by glutamate or GABA, we conducted post hoc analyses of these associations. Here we found increasing response latency was marginally associated with increasing glutamate (*β* = 0.75, t = 2.50, *p* = 0.05; Figure 11B), but no association with GABA (*β* = −0.05, t = - 0.34, *p =* 1.00; Figure 11C).

**Figure 11.**
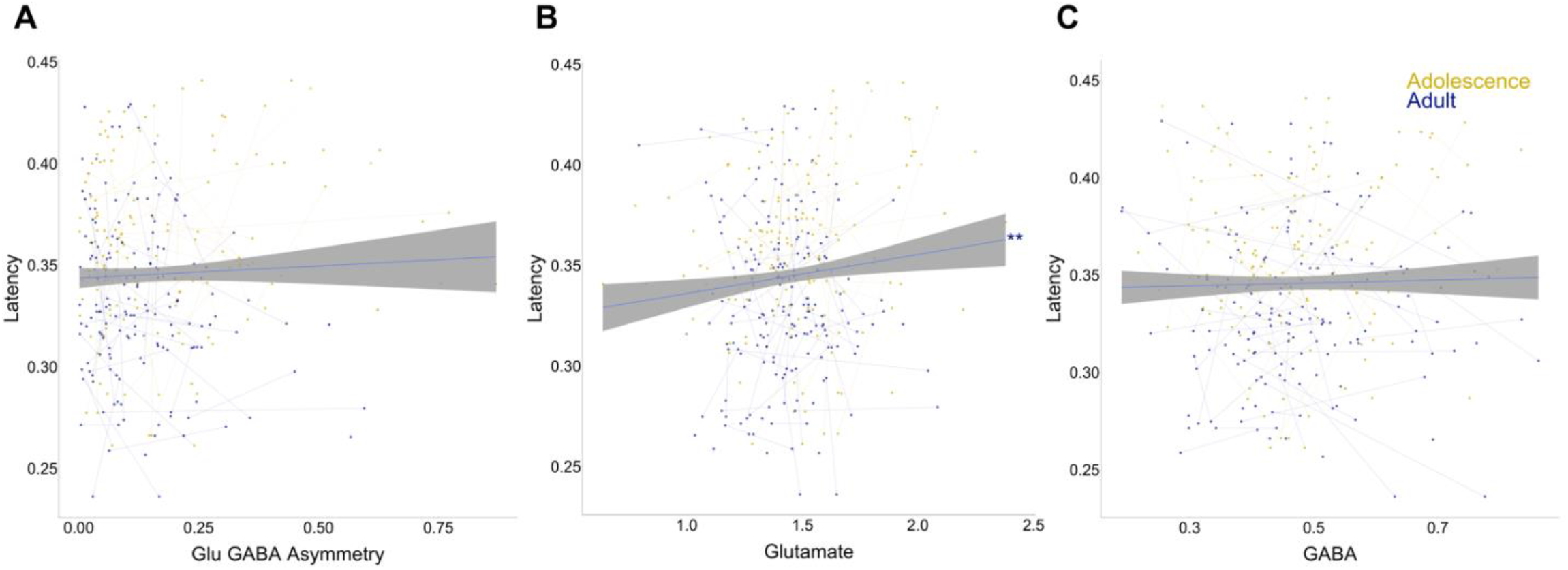
MRSI associations with behavioral measures. Adolescence is all participants under 18 (gold) and adult represents all participants 18 and above (purple). **A**. Glu GABA Asymmetry vs MGS response latency **B**. Glu vs MGS response latency **C**. GABA vs response latency (* p < 0.01)

## 4. DISCUSSION

Using novel EEG and MRSI methodology in a large, longitudinal, developmental dataset through adolescence, this study demonstrates an association between developmental changes in DLPFC glutamate and GABA and changes in DLPFC EEG markers of neural excitatory and inhibitory activity. As hypothesized, we found age-related decreases in both the offset and exponent parameters of EEG-based aperiodic activity across conditions, as previously reported^20,41–43^. Furthermore, in accordance with our prior findings, we found that balance between glutamate and GABA increased through adolescence. Post hoc analyses highlighted the developmental changes we see in glutamate GABA asymmetry may be driven by significant age-related decreases in glutamate, as no significant changes in GABA across adolescence have been found. Importantly, we found that glutamate/GABA asymmetry is associated with EEG-derived exponent but not offset, with more asymmetry between glutamate and GABA being associated with larger exponent. However, glutamate was found to significantly decrease with decreasing offset. Furthermore, we found that glutamate/GABA asymmetry mediates the association between age and exponent, while glutamate mediates the associations between age and offset, providing a neurobiological mechanistic link between age and aperiodic neural activity. Upon further interrogation into the developmental effects the E/I balance may have on behavior, we assessed working memory via two working memory tasks: the memory guided saccade (MGS) task and the CANTAB Spatial Span task. In line with prior findings^57^, we observed robust developmental improvements in working memory that were significantly associated with decreases in glutamate as well as decreases in the aperiodic offset. The spatial span measure, indicating the maximum length of a memorized sequence, was found to improve with decreasing offset and exponent. Taken together, these results suggest neurobiological mechanisms of excitation and inhibition, measurable through EEG and MRSI, may be associated with the development of higher order cognitive functions across adolescence.

Previous work has stressed the importance of interpreting neurophysiological recordings as a combination of periodic (oscillatory) and aperiodic components. Age-related differences in band specific oscillations may thus arise in part due to a shift in the aperiodic slope (exponent) or intercept (offset)^24^. Our findings show that exponent and offset decrease with age during adolescence, largely in line with the extant literature, including prior findings of flatter 1/f slopes (exponent) and reduced offset values in older children compared to younger children^20,41–43^. This result has been extended in adult populations, with exponent and offset decreasing across the lifespan^37,58^. These results have been interpreted based on the hypothesized neurophysiological underpinnings of these measures. The spectral slope is thought to reflect the E/I balance via decays in local field potential (LFP) power constrained by faster decaying excitatory AMPA and slowly decaying inhibitory GABA currents^31^, with steeper slopes representing inhibitory activity dominating over excitatory^24^. In parallel, the amplitude of the spectra, assessed by the aperiodic offset parameter, is associated with neural spike rates^59,60^, supported by intercranial LFP recordings that have shown increases in broadband LFP power associated with neuronal population spiking^22^. Shifts in excitatory-inhibitory circuits, reflected by a change in the aperiodic model parameters, may thus reflect cortical maturation including refinement of cortical networks, synaptic pruning, myelination, and alterations in the E/I balance^20,21,61^. Thus, our results showing robust reductions in aperiodic offset could be indicative of maturational decreases in overall spike rate of cortical neurons, while our results showing decreases in exponent suggest maturational increases in the E/I balance.

In our previous work in a cross-sectional study we found increasing balance between glutamate and GABA across frontal regions, including DLPFC, across adolescence into adulthood^18^. Here, we confirm these results but now in a longitudinal sample. Structurally, excitatory and inhibitory synapses maintain a fairly constant ratio^63^, the location of which are regulated by changes in connectivity as a result of synaptic pruning^64^. Functionally, the E/I currents maintain balance via a “push-pull” mechanism, constantly blocking and activating excitatory and/or inhibitory receptors^64^. Mice hippocampal cultures and modeling work has shown that for networks with a range of E/I fractions, ranging from 10 to 80% inhibitory neurons, the networks maintain stable neuronal activity^64^. Developmental changes in E/I balance may reflect an increase in “optimal” quantity of E and I activity that provides a state of low firing rates without hyper or hypo activity in response to stimuli. Developmentally adulthood is the “optimal” network state, thus, age-related decreases in the Glu GABA asymmetry across adolescence may be reflecting the optimization of the excitatory and inhibitory currents that result in balance. Furthermore, post-hoc analyses revealed glutamate to be significantly decreasing across adolescence. Levels of Glu/Cr as measured by spectroscopy have been linked to PET-derived measures of synaptic density^65^. During adolescent development, synaptic pruning of primarily excitatory synapses has been shown to take place in prefrontal cortex^66^, thus age-related pruning of synapses could lead to decreases in levels of glutamate and decreased neural population activity as indexed by offset. Future studies should aim to measure synaptic pruning directly to establish this link to modulation of excitatory activity and E/I balance.

Importantly, we found associations between the aperiodic exponent and Glu GABA asymmetry with decreasing asymmetry being significantly associated with decreasing exponent. Furthermore, we found that age-related changes in exponent are mediated by Glu GABA asymmetry reflecting how correlated excitation and inhibition are to one another. This is consistent with prior work showing that modulating the E/I balance in either direction (increasing E relative to I or increasing I relative to E) via pharmacological manipulation produced similar changes in aperiodic components (i.e., steepening of the slopes)^67,68^. Consistent with this literature, we showed that decreases in Glu GABA asymmetry led to flattening of the aperiodic slope. This result suggests that the Glu GABA asymmetry measure may be more indicative of the E/I balance, via the changes in correlation between excitatory and inhibitory neurons^69^ and may support the notion that E/I balance may be inferred through the PSD slope exponent, providing important methodological backing to the FOOOF protocol using empirical longitudinal data and vice versa. In addition, post-hoc analyses revealed associations between Glu/Cr and aperiodic offset as well as exponent, with lower Glu/Cr levels associated with decreased offset and exponent. This suggests that age-related pruning of synapses may contribute to decreases in levels of glutamate and decreased neural population activity as indexed by offset.

Using two working memory measures, the MGS, and the Spatial Span Task, we assessed whether working memory was associated with our measures of E/I balance. DLPFC is a region that is known to be involved in visuo-spatial working memory^70^, and exhibits a protracted developmental trajectory through adolescence^71^. Both accuracy and latency, derived from the MGS task, are important but distinct components underlying overall executive function, with accuracy reflecting number of correct responses and latency reflecting the speed of information processing prior to making a response^8,72^ Using the MGS task, we found significant associations between decreasing offset and decreasing response latency. While we did not find any associations between Glu GABA asymmetry and accuracy or latency, post hoc analyses showed a significant negative relationship between response latency and glutamate. These results suggest that the age-related decrease in neural population spiking, reflected by the decreased offset and decreased glutamate, may contribute to decreased response latency, suggesting adults may need less excitatory neuronal activity in order to perform the task quickly. Similarly, the association between spatial span and decreases in exponent suggest that age related improvement in working memory capacity are supported by the optimization of the balance of excitation and inhibition^73–75^.

Together, these findings provide novel *in vivo* evidence describing neurobiological mechanisms underlying the maturation of PFC neural function and cognition throughout adolescence. Importantly, these findings are in line with a model of adolescence as a critical period for frontal cortex supporting cognitive development into adulthood^10^. In this model, developmental changes in glutamate and GABA would drive changes in the E/I balance, supporting maturation of neural activity that supports optimal cognitive control. Developmental disruptions in the development of E/I balance in frontal cortex have been associated with cognitive deficits associated with psychopathology, such as schizophrenia^76,77^. Understanding normative PFC development and plasticity can inform mechanisms underlying deviations from normative trajectories and possible interventions.

## Acknowledgements

We thank the University of Pittsburgh Clinical and Translational Science Institute (CTSI) for support in recruiting participants, as well as their support by the National Institutes of Health through grant number UL1TR001857. We thank Matthew Missar, Laurie Thompson, Alyssa Famalette, and Vivian Lallo for their work involving our data collection.

## Notes

**Funding:** This work was supported by MH067924 from the National Institute of Mental Health, T32 Training Grant Number T32MH019986 from the National Institute of Mental Health, the Staunton Farm Foundation, and support from the Department of Bioengineering, University of Pittsburgh.

**Declaration of Interests:** The authors declare no conflict of interests.

### Competing Interest Statement

The authors have declared no competing interest.

