## Supplemental for "Aperiodic EEG and 7T MRSI evidence for maturation of E/I balance supporting the development of working memory through adolescence"

**Supplementary Figures and Tables**

*Model 1: Aperiodic parameter ~ s(age, k = 3, fx = T) + hemisphere + condition, random = list(subject =~1)*

*Model 2: Aperiodic parameter ~ s(age, k = 3, fx = T) + s(age, by = hemisphere k = 3, fx = T) + condition, random = list(subject =~1)*

*Model 3: Aperiodic parameter ~ s(age, k = 3, fx = T) + s(age, by = condition k = 3, fx = T) + hemisphere, random = list(subject =~1)*

*Model 4: MRS measure ~ s(age, k = 3, fx = T) + fracGM + hemisphere, random = list(subject =~1)*

*Model 5: MRS measure ~ s(age, k = 3, fx = T) + s(age, by = hemisphere k = 3, fx = T) + fracGM, random = list(subject =~1)*

*Model 6: Aperiodic parameter ~ MRS measure + inverse age + hemisphere + condition + (1 | Subject)*

*Model 7: Aperiodic parameter ~ MRS measure*inverse age + hemisphere + condition + (1 | Subject)*

*Model 8: Aperiodic parameter ~ Behavioral measure + inverse age + hemisphere + condition + (1 | Subject)*

*Model 9: Aperiodic parameter ~ Behavioral measure*inverse age + hemisphere + condition + (1 | Subject)*

*Model 10: MRS Measure ~ Behavioral measure + age (or inverse age) + hemisphere + (1 | Subject)*

*Model 11: MRS Measure ~ Behavioral measure*age (or inverse age) + hemisphere + (1 | Subject)*

Table S1. AIC values for FOOOF measures vs MRS measures

| | Exponent | | Offset | |
|---|---|---|---|---|
| | Linear AIC | Inverse AIC | Linear AIC | Inverse AIC |
| Glu | -389.59 | **-418.60** | -278.95 | **-349.03** |
| GABA | -351.20 | **-379.51** | -233.04 | **-301.53** |
| Imbalance | -353.17 | **-377.89** | -250.41 | **-312.71** |

Table S2. AIC values comparing linear and inverse models for all associations between FOOOF and behavior and MRS measures and behavior

| | Accuracy | | Accuracy Var | | Latency | | Latency Var | | Spatial Span Max | |
|---|---|---|---|---|---|---|---|---|---|---|
| | Linear AIC | Inverse AIC | Linear AIC | Inverse AIC | Linear AIC | Inverse AIC | Linear AIC | Inverse AIC | Linear AIC | Inverse AIC |
| Exponent | -356.28 | **-393.16** | -352.39 | **-388.64** | -370.01 | **-408.73** | -373.67 | **-411.38** | -302.43 | **-319.23** |
| Offset | -237.45 | **-317.29** | -240.66 | **-316.90** | -252.77 | **-336.78** | -247.50 | **-326.33** | -218.93 | **-245.19** |
| Glu | -163.55 | **-175.14** | -144.81 | **-156.64** | -135.86 | **-147.48** | -153.60 | **-165.76** | -222.20 | **-234.94** |
| GABA | -886.30 | **-899.90** | -890.01 | **-904.51** | -869.61 | **-881.73** | -897.52 | **-911.02** | -656.65 | **-668.41** |
| Imbalance | -726.60 | **-742.58** | -724.96 | **-742.40** | -692.73 | **-708.38** | -734.28 | **-750.68** | **1029.15** | 1015.99 |

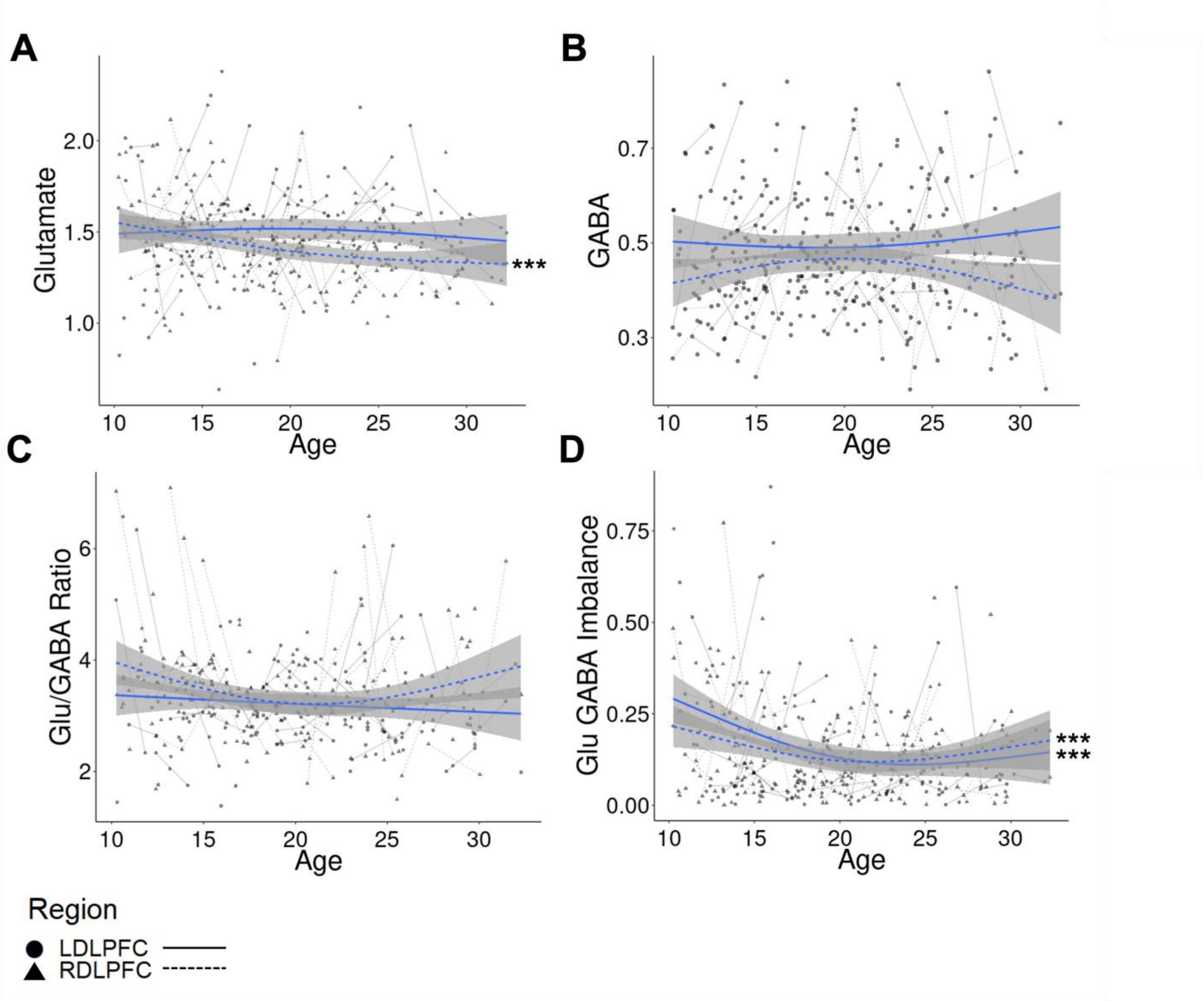


Figure S1. **A.** Glutamate vs age had main effects of hemisphere (F = 15.25, p < 0.0001), and an age-by-hemisphere interaction (F = 3.46, p = 0.03), where follow-up tests revealed that the effect of age was stronger in the right hemisphere (F = 13.12, p = 0.0004) in comparison to the left hemisphere (F = 0.44, p = 0.51). **B.** GABA vs age had a main effect of hemisphere (left vs right DLPFC; F = 15.72, p < 0.0001), but no age-by-hemisphere interactions (F = 2.6, p = 0.08) suggesting uniform effects of age across both hemispheres (Left DLPFC: F = 0.26, p = 0.612; Right DLPFC: F = 1.11, p = 0.266). **C.** The Glu/GABA ratio had a main effect of hemisphere (F = 5.24, p = 0.02), but no age-by-hemisphere interaction (F = 2.27, p = 0.11), suggesting similar effects of age across both hemispheres (Left DLPFC: F = 2.27, p = 0.133; Right DLPFC: F = 4.2, p = 0.04). **D.** The Glu GABA imbalance measure had no main effect of hemisphere (F = 0.95, p = 0.33), and no significant age-by-hemisphere interaction (F = 1.85, p = 0.16). Thus, both hemispheres have similar age-related changes (Left DLPFC: F = 12.57, p = 0.0006; Right DLPFC: F = 2.7, p = 0.18).

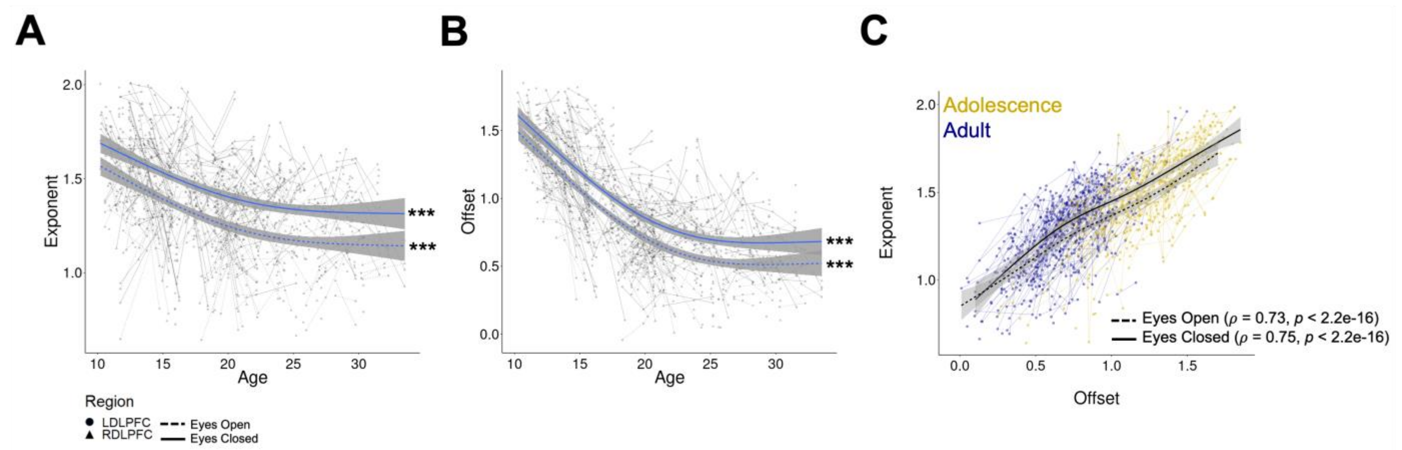


Figure 2. Aperiodic activity of EEG during resting state. **A.** Exponent across adolescence across both eyes closed (solid line) and eyes open (dashed line) conditions. The exponent had a main effect of condition (F = 275.22, p < 0.001), and a significant age-by-condition (eyes open or eyes closed) interaction (F = 3.31, p = 0.02) reflecting greater age-related decreases during eyes open (F = 50.17, p < 0.0001) compared to eyes closed (F = 49.2, p < 0.0001). Further, there was no main effect of hemisphere (F = 1.37, p = 0.24), nor age-by-hemisphere interactions (F = 0.61, p = 0.61). **B.** Offset across adolescence across both eyes closed (solid line) and eyes open (dashed line). Similar to the exponent parameter, there was a significant main effect of condition (F = 247.97, p < 0.0001), indicating lower power during eyes open compared to eyes closed, with an age-by-condition interaction (F = 4.53, p = < 0.0001), reflecting greater age-related decreases during eyes open (F = 50.17, p < 2e-16) compared to eyes closed (F = 49.22, p < 2e-16). There was a main effect of hemisphere (F = 7.66, p = 0.01), but no age-by-hemisphere interaction (F = 0.75, p = 0.52). **C.** Correlation between exponent and offset values for both the left and right DLPFC for both conditions.

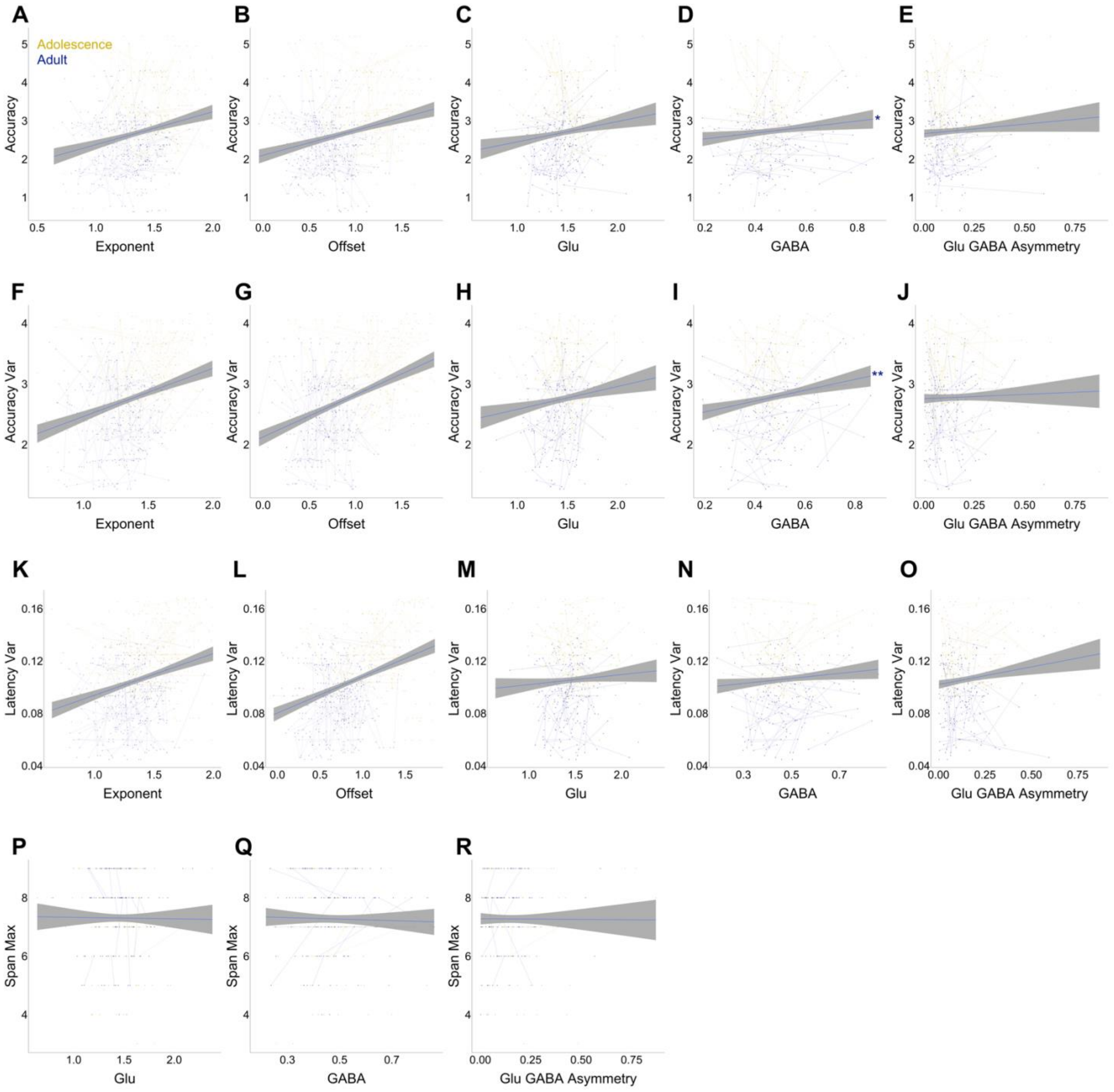


Figure S3. Associations between behavioral measures and 1/f aperiodic activity and MRS measures. **A.** Exponent vs accuracy **B.** Offset vs accuracy **C.** Glu vs accuracy **D.** GABA vs accuracy **E.** Glu GABA Asymmetry vs accuracy **F.** Exponent vs accuracy variability **G.** Offset vs accuracy variability **H.** Glu vs accuracy variability **I.** GABA vs accuracy variability **J.** Glu GABA Asymmetry vs accuracy variability **K.** Exponent vs Latency variability **L.** Offset vs Latency variability **M.** Glu vs Latency variability **N.** GABA vs Latency variability **O.** Glu GABA Asymmetry vs Latency variability **P.** Glu vs Spatial Span Max **Q.** GABA vs Spatial Span Max **R.** Glu GABA Asymmetry vs Spatial Span Max

Table S3. Associations between behavior and FOOOF parameters, exponent and offset, and MRS measure, as. All p values are Bonferroni corrected.

| | | Exponent | | | Offset | | | Glu | | | GABA | | | Asymmetry | | |
|---|---|---|---|---|---|---|---|---|---|---|---|---|---|---|---|---|
| | | **β** | **t** | **p** | **β** | **t** | **p** | **β** | **t** | **p** | **β** | **t** | **p** | **β** | **t** | **p** |
| **Behavioral main effects** | **Accuracy** | -0.01 | -1.16 | 0.99 | -0.01 | -1.04 | 1.00 | 0.02 | 1.66 | 0.39 | 0.02 | 2.55 | **0.04** | -0.01 | -0.70 | 1.00 |
| | **Accuracy Var** | 0.00 | 0.06 | 1.00 | 0.04 | 2.05 | 0.16 | 0.03 | 1.36 | 0.69 | 0.04 | 3.85 | **0.00** | -0.03 | -2.22 | 0.11 |
| | **Latency** | -0.63 | -2.44 | 0.06 | -1.04 | -3.60 | **0.00** | 0.75 | 2.50 | **0.05** | -0.05 | -0.34 | 1.00 | -0.06 | -0.31 | 1.00 |
| | **Latency. Var** | -0.32 | -0.69 | 1.00 | -0.71 | -1.34 | 0.73 | 0.04 | 0.08 | 1.00 | 0.60 | 2.14 | 0.13 | -0.12 | -0.35 | 1.00 |
| | **Spatial Span Max** | -0.04 | -3.09 | **0.01** | -0.04 | -3.12 | **0.01** | .003 | 0.23 | 1.00 | .003 | 0.44 | 1.00 | 0.01 | 0.95 | 1.00 |
| **Behavioral age interaction** | | **β** | **t** | **p** | **β** | **t** | **p** | **β** | **t** | **p** | **β** | **t** | **p** | **β** | **t** | **p** |
| | **Accuracy** | 1.62 | 1.92 | 0.22 | 1.87 | 1.95 | 0.21 | -1.41 | -1.60 | 0.11 | -0.28 | -0.59 | 0.56 | 0.35 | 0.58 | 1.00 |
| | **Accuracy Var** | 0.73 | 0.66 | 1.00 | 1.06 | 0.88 | 1.00 | -2.75 | -2.29 | 0.02 | -1.31 | -2.06 | 0.04 | -0.58 | -0.74 | 1.00 |
| | **Latency** | 25.6 | 1.72 | 0.35 | 23.9 | 1.29 | 0.80 | 40.3 | 2.40 | 0.02 | 8.97 | 1.02 | 0.31 | 32.3 | 3.01 | **0.01** |
| | **Latency. Var** | 44.1 | 1.56 | 0.48 | 14.2 | 0.41 | 1.00 | -2.01 | -0.07 | 0.95 | -7.20 | -0.47 | 0.64 | 43.2 | 2.22 | 0.11 |
| | **Spatial Span Max** | 0.17 | 0.26 | 1.00 | 0.05 | 0.64 | 1.00 | -0.37 | -0.54 | 1.00 | -0.54 | -1.58 | 0.47 | 0.25 | 0.55 | 1.00 |
